# Tannic cell walls form a continuous apoplastic barrier sustaining Arabidopsis seed coat biophysical properties

**DOI:** 10.1101/2020.05.14.096669

**Authors:** Lara Demonsais, Anne Utz-Pugin, Sylvain Loubéry, Luis Lopez-Molina

## Abstract

Seeds are a late land plant evolution innovation that promoted the striking spread and diversity of angiosperms. The seed coat is a specialized dead tissue protecting the plant embryo from mechanical damage. In many species, including *Arabidopsis thaliana*, the seed coat also achieves a remarkable balancing act: it limits oxygen uptake, avoiding premature embryo oxidative damage, but not entirely so as to enable seed dormancy release. The seed coat biophysical features implementing the striking physiological properties of the seed remain poorly understood. Tannins, a type of flavonoids, are antioxidants known to accumulate in the Arabidopsis seed coat and *transparent testa* (*tt*) mutant seeds, deficient in flavonoid synthesis, exhibit low dormancy and viability. However, their precise contribution to seed coat architecture and biophysics remains evasive. A seed coat cuticle, covering the endosperm outer surface was, intriguingly, previously shown to be more permeable in *tt* mutants deficient not in cuticular component synthesis, but rather in flavonoid synthesis. Investigating the role of flavonoids in cuticle permeability led us to identify cell walls, originating from the seed coat inner integument 1 cells, impregnated with tannins. We found that tannic cell walls are tightly associated with the cuticle, forming two fused layers that regulate endosperm permeability. In addition, we show that tannic cell walls are prominent building blocks of the seed coat, constituting a continuous barrier around the seed living tissues. Altogether our findings reveal the existence of tannic cell walls as a previously unrecognized biological barrier sustaining the seed’s key physiological properties.

**One sentence summary:** The seed coat is largely composed of plant cell walls impregnated with tannins, forming a thick and continuous protective barrier surrounding the embryo promoting seed viability and dormancy.

## INTRODUCTION

Land plant ancestors were aquatic organisms that evolved to cope with a water-deprived environment (Cheng et al., 2019a). Seeds are a late land plant evolution innovation consisting of plant embryos maintained in a desiccated and highly sheltered state surrounded by a protective dead seed coat of maternal origin. By enhancing plant embryo survival and dispersion, seeds undoubtedly contributed to the spectacular spread of angiosperms as they virtually converted plants into time and space travelers. However, in contrast to the plant’s vegetative phase, where indeterminate development facilitates adaptation to an ever-changing environment, embryo survival only relies on the pre-established highly resistant mature seed state that notably depends on the protective seed coat. Although seeds are desiccated and metabolically inert structures, their exposure to atmospheric oxidation will irremediably lead to their oxidative damage. Besides its obvious mechanical protective function, the seed coat fulfills the need to shield the living tissues from excessive exposure to atmospheric oxygen and is obligatory in order to preserve the long-term capacity of the seed to produce a viable seedling. Furthermore, the seed coat also regulates the entry of water upon seed imbibition, potentially to discriminate partial or short-term water exposure from more prolonged imbibition (Windsor et al., 2000; De Giorgi et al., 2015). Hence, understanding the biophysical and anatomical properties of the boundary demarcating the seed’s living tissues from the environment is essential to understand seed physiology and more generally plant fitness.

The model organism *Arabidopsis thaliana* has revealed what sort of seed coat characteristics can contribute to the remarkable physiological properties of seeds. In Arabidopsis, the mature seed consists of an external dead seed coat surrounding a single cell layer of endosperm itself surrounding the embryo. The seed coat arises from the differentiation of ovular integuments after double fertilization that produces the endosperm and zygote. Seed germination takes place upon seed imbibition and is defined by concomitant embryonic radicle emergence and endosperm rupture. However, seed coat rupture is the first visible event prior to seed germination and therefore defines the time whereupon the endosperm is exposed to the outer environment. During ovular differentiation accompanying seed development, three cell layers of inner integuments (ii1, ii1’, ii2) and two cell layers of outer integuments (oi1 and oi2) undergo complex modifications, giving rise to the mature seed coat (Fig. 1A). Inner and outer ovular integuments will contribute distinct key components of the mature seed coat such as production of tannins and mucilage, respectively. The seed coat brown pigment layer (bpl) arises from the progressive collapse of ii1’ and ii2 cells during late seed development (Fig. 1A) (Beeckman et al., 2000; Debeaujon et al., 2007).

**Figure 1.**
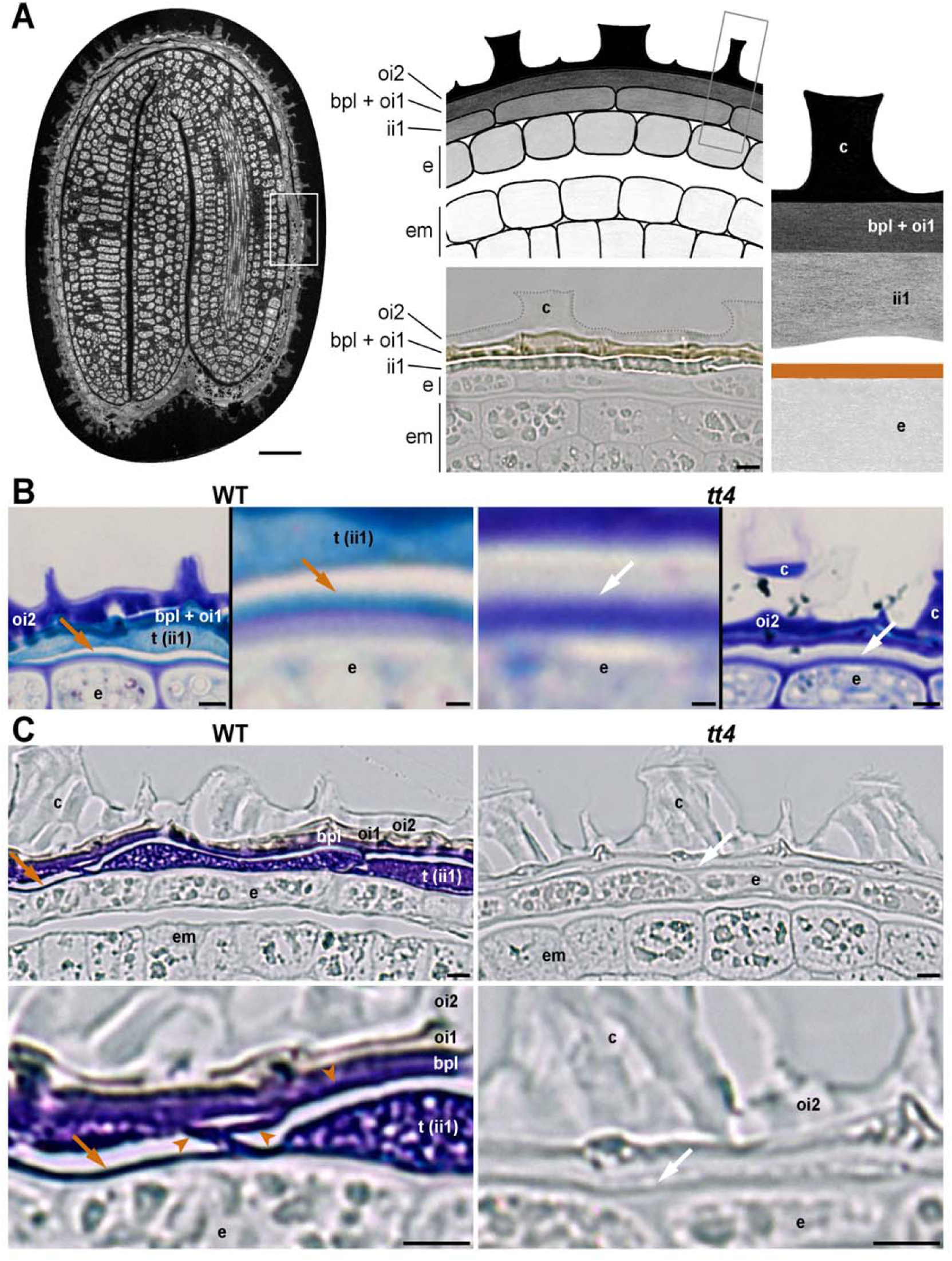
Presence of tannins at the outer surface of the endosperm. (A) Left, semi-thin section of a WT seed (autofluorescence). Center top, schematic drawing illustrating the organization of the seed peripheral area (corresponding to the white-boxed region in the left panel). Center bottom, micrograph of an unstained semi-thin section showing the seed peripheral area; ii1, the bpl, oi1 and oi2 form the seed coat. Right panel, schematic drawing of an enlarged seed coat area (grey-boxed region in the center top panel), the brown line indicates the localization of tannins along the endosperm outer surface (arrows in B-C and in Supplemental Figure S1, A-C). Drawings were not made to scale in order to better visualize the positions of the different cells and structures of interest. (B) Semi-thin sections of WT seeds (left panels) and *tt4* seeds (right panels) stained with toluidine blue; in each case, a global view of the seed coat and an enlargement of the region around the arrow is shown. (C) Semi-thin sections of WT seeds (left panels) and *tt4* seeds (right panels) stained with DMACA; in each case, a global view of the seed coat and an enlargement of the region around the arrow is shown below. In B and C, brown arrows (left panels) indicate the detection of tannins along the endosperm outer cell wall; white arrows (right panels) indicate no detection of tannins along this cell wall. Brown arrowheads in C (bottom left panel) indicate the reticulated structure formed by ii1 tannic cell walls. Scale bars: A, 50 μm (left) and 5 μm (center bottom); B, 5 μm and 1 μm for general views and magnified insets, respectively; C, 5 μm. Abbreviations: bpl, brown pigment layer; c, columella; e, endosperm; em, embryo; ii1, inner integument 1; oi1, outer integument 1; oi2, outer integument 2; t, tannins.

At early seed developmental stages, ii1 cells, differentiating alongside the developing endosperm, begin to accumulate proanthocyanidins, a type of oligomeric flavonoids also known as condensed tannins, and hereafter referred to as tannins. Oxidation of tannins in the ii1 layer is often proposed to account for the characteristic brown color of WT Arabidopsis mature seeds. Juxtaposed to the ii1 layer, the bpl remains poorly characterised, in particular regarding its composition. It has been suggested that the bpl contains phenolic compounds, which were proposed to also participate in the final mature seed color (Beeckman et al., 2000; Debeaujon et al., 2000, 2001).

Debeaujon and Kornneef have demonstrated the importance of tannins in seed coat physiology. Indeed, a suite of *transparent testa* (*tt*) mutants, deficient in flavonoid synthesis and producing colorless seeds, have low dormancy and viability as well as premature seed coat rupture (Debeaujon et al., 2000). *tt* mutants exhibit enhanced permeability to the dye tetrazolium red and, accordingly, these defects can be associated with increased oxidative damage (Landry et al., 1995; Debeaujon et al., 2000; MacGregor et al., 2015). Importantly, *tt* mutant phenotypes are maternally inherited and therefore the phenotypes are due to a deficient seed coat. A number of explanations have been put forward regarding the function of flavonoids in seeds. Flavonoids act as antioxidants and therefore their absence could explain the increased oxidative damage observed in *tt* mutant seeds. Furthermore, it was hypothesized that during seed desiccation, oxidation of phenolic compounds, such as tannins, would reduce seed coat permeability (Marbach and Mayer, 1974; Stafford, 1974; Marbach and Mayer, 1975; Werker et al., 1979). However, tannins have been described as filling the intracellular spaces of ii1 cells: in this view, they form spatially separated discrete units, separated by cell walls (Bowman, 2011). How such a discontinuous arrangement would limit oxygen diffusion and influence seed coat rupture is unclear. More generally, how tannins participate in the seed coat permeability is not well understood.

The cuticle is an evolutionary innovation that enabled early land plant colonizers to conserve water, essential for cellular function, in a water-limited terrestrial environment. Indeed, cuticles are fatty and waxy hydrophobic films covering the plant surface (Yeats and Rose, 2013). Recently, a cuticle tightly associated with the outer cell wall of the endosperm in mature seeds was identified (De Giorgi et al., 2015). Remarkably, this cuticle is maternal, being produced by ii1 cells during seed development and transferred to the endosperm around the heart and torpedo stages of embryo development (Loubéry et al., 2018). Genetic studies showed that mutants defective in cuticle biosynthesis have low viability and dormancy levels, which correlates with increased oxidative stress in mature seeds (De Giorgi et al., 2015). In addition, similar to *tt* mutants, cuticle biosynthesis mutants prematurely ruptured their seed coat upon seed imbibition as a result of abnormal endosperm cell expansion (De Giorgi et al., 2015). Furthermore, in seed coat-ruptured seeds, the endosperm of cuticle biosynthesis mutants was shown to be more permeable to toluidine blue, a dye commonly used to assess cuticle integrity (Tanaka et al., 2004; De Giorgi et al., 2015). These observations are consistent with the notion that a deficient endosperm-associated cuticle disrupts the seed’s water uptake dynamics and increases the seed’s permeability to oxygen. They also indicate that during evolution plants recycled an old structure, the cuticle, to achieve the biophysical properties of the seed coat.

However, a number of unsolved questions regarding the endosperm-associated cuticle remain unaddressed. Firstly, whether the maternal cuticle continues to be associated with the endosperm upon seed imbibition and therefore whether it directly contributes to its permeability was not investigated. Secondly, Loubéry et al. reported that the endosperm of *tt* mutant seeds also showed increased toluidine blue permeability, suggesting that the endosperm-associated cuticle is deficient in *tt* mutants (Loubéry et al., 2018). How a deficiency in synthesis of flavonoids impacts the permeability of the endosperm-associated cuticle was not investigated. Histological experiments suggested that tannins were present on the outer surface of the endosperm (Loubéry et al., 2018). However, whether their presence simply reflects nonspecific binding to the endosperm outer surface following their release after seed coat rupture was not investigated. Alternatively, whether tannins are *bona fide* components of the endosperm-associated cuticle is unknown. Hence, whether tannins directly or indirectly participate in the permeability of the cuticle remains to be understood. Addressing this specific question would provide insights about how tannins generally contribute to the seed coat biophysical properties.

Here we show that the endosperm-associated cuticle is part of a previously uncharacterized seed structure. The outer surface of the endosperm is covered by a bilayer of electron-lucent and electron-dense material (De Giorgi et al., 2015). We show that the former corresponds to the cuticle, whereas the latter is a newly described structure in seeds: a cell wall impregnated with tannins, which originates from ii1 cells and referred to here as tannic cell wall. We show that tannic cell walls remain attached to the endosperm upon seed coat rupture, thus participating in its permeability properties. Furthermore, tannic cell walls leave a characteristic imprint of ii1 cell wall material on the outer endosperm surface; this reveals that during seed coat rupture, the ii1 cell walls anticlinal to the endosperm surface are breaking points between the inner seed tissues and the seed coat. Lastly, our study reveals that tannic cell walls are not restricted to the inner side of ii1 cells. We describe the existence of an extended and continuous network of tannic cell walls, which includes the ii1 and the bpl cell layers, providing a continuous seal shielding the living tissues from water and atmospheric gases. Collectively, our findings shed new light on how the biophysical properties of the Arabidopsis seed coat are implemented by means of a cuticle and tannic cell walls.

## RESULTS

It was shown using *transparent testa* (*tt*) mutant seeds, deficient in flavonoid synthesis, that flavonoids are necessary to confer normal impermeability to the outer surface of the endosperm and, by extension, to the endosperm-associated cuticle (Loubéry et al., 2018). However, whether the involvement of flavonoids is direct, i.e. as intrinsic components of the cuticle, or indirect was not determined. Furthermore, whether the endosperm-associated cuticle remains attached or not at the surface of the endosperm upon rupture of the seed coat was not ascertained. Here we sought to address these two questions in turn.

### Presence of tannins at the outer surface of the endosperm

Histological sections by Loubéry et al. suggested that the surface of mature endosperm cells contains tannins, since it appeared brown in WT seeds but not in *transparent testa 4* (*tt4*) mutants, lacking the enzyme for the first step of flavonoid biosynthesis and hence lacking tannins (Feinbaum and Ausubel, 1988; Loubéry et al., 2018). However, the resolution of the histological preparations did not allow identifying unambiguously the subcellular structure harboring this color, i.e. to discriminate whether it was the cuticle itself that was brown or another closely apposed structure. To further explore these possibilities we sought to clarify the putative presence of tannins along the endosperm surface (brown line in Fig. 1A, right), using a combination of genetics and of improved histological methods.

We used higher resolution semi-thin sections of Epon-embedded mature dry seeds stained with toluidine blue and methylene blue. These two metachromatic dyes revealed tannins in the ii1 cell layer with blue (toluidine blue) and turquoise (methylene blue) colors (Fig. 1B; Supplemental Fig. S1A, left panels). Strikingly, they also revealed a linear structure along the outer surface of the endosperm that shared the same color as that of tannins (Fig. 1B; Supplemental Fig. S1A, left panels). The similarity in color obtained with two different metachromatic dyes suggests a similarity in composition: thus, these data suggest that the linear structure might contain tannins. Consistently, these linear structures were not colored in sections of *tt4* mature seeds, lacking tannins, upon staining with the same dyes (Fig. 1B; Supplemental Fig. S1A, right panels). Similarly, in unstained WT seed sections, a dark beige line was clearly visible along the endosperm surface that had the same color and shade as that of the tannins present in the ii1 cell layer (Supplemental Fig. S1B, left panels). By changing the focal plane, the line showed a yellowish color, which was similar to that shown by the tannins (Supplemental Fig. S1C, left panels), and as above, it was not visible in *tt4* mutants (Supplemental Fig. S1, B and C).

4-dimethylaminocinnamaldehyde (DMACA) is a dye that binds tannins and its late precursors (flavan-3-ols) with high specificity, as it does not bind other flavonoids such as flavonols or anthocyanidins (Cadot et al., 2006; Auger et al., 2010; Hammouda et al., 2014). DMACA presents a significantly better sensitivity than vanillin, another dye commonly employed to detect tannins (Li et al., 1996); besides, its purple/deep blue signal is best-suited for Arabidopsis seed coat histological studies, compared with vanillin whose light orange/pink signal can easily be confused with the natural brownish color of the seed coat (Li et al., 1996; Abrahams et al., 2002; Kitamura et al., 2004). Upon DMACA staining of WT semi-thin sections, tannins in ii1 cells gave a purple signal; in addition, a purple signal was visible on a linear structure along the outer surface of the endosperm (Fig. 1C, left panels). As above, this linear signal was not visible in *tt4* mutant semi-thin sections (Fig. 1C, right panels).

Altogether, these results demonstrate the presence of tannins at the outer surface of endosperm cells in Arabidopsis mature seeds. Next, we sought to identify which structures at the outer surface of the endosperm contain tannins.

### ii1 cell walls are tannified and form a continuous reticulated tannic apoplastic barrier tightly bound to the endosperm-associated cuticle

In previous reports, transmission electron microscopy (TEM) images revealed the presence of a two-layered structure covering the outer surface of mature endosperm cells, consisting of an inner electron-lucent layer and an outer electron-dense layer (De Giorgi et al., 2015; Loubéry et al., 2018). The two layers were interpreted as representing different compositions of the endosperm-associated cuticle. Furthermore, the electron-dense layer periodically protruded outwardly, forming an extended electron-dense reticulated structure surrounding the endosperm (purple dashed line in Supplemental Figure S2). Loubéry et al. proposed that this reticulated structure is an apoplastic network consisting of electron-dense cuticular material enclosing ii1 cells. However, Loubéry et al. noted that this is inconsistent with the fact that this structure was not recognized by the cuticle dye Auramine O (Loubéry et al., 2018). The nature and the composition of these layers and that of the reticulated structure was not further investigated.

Here, we challenge the notion that the electron-dense and electron-lucent layers represent different compositions of a cuticle. Rather, we propose that only the electron-lucent layer is a *bona fide* cuticle; we also propose that the electron-dense layer corresponds to ii1 cell walls that have undergone tannin impregnation (Fig. 2A). Hereafter, we refer to tannin-impregnated cell walls as tannic cell walls. This model therefore states that during seed development, the ii1 cells produce a cuticle on their inner side, while their primary cell walls become impregnated with tannins. Eventually, the inner cuticle and inner tannified cell wall become tightly associated with the outer surface of the endosperm (red-boxed region in Fig. 2A). The remaining ii1 tannified walls form the reticulated structure described by Loubéry et al., which generates a continuous tannic barrier (Fig. 2A; Supplemental Fig. S2). In turn, in each ii1 cell, the tannic cell walls enclose the tannic depositions that are visible as rugged electron-dense blocks in TEM. The tannic cell wall described by this model represents, to our best knowledge, a previously uncharacterized structure in Arabidopsis seeds.

**Figure 2.**
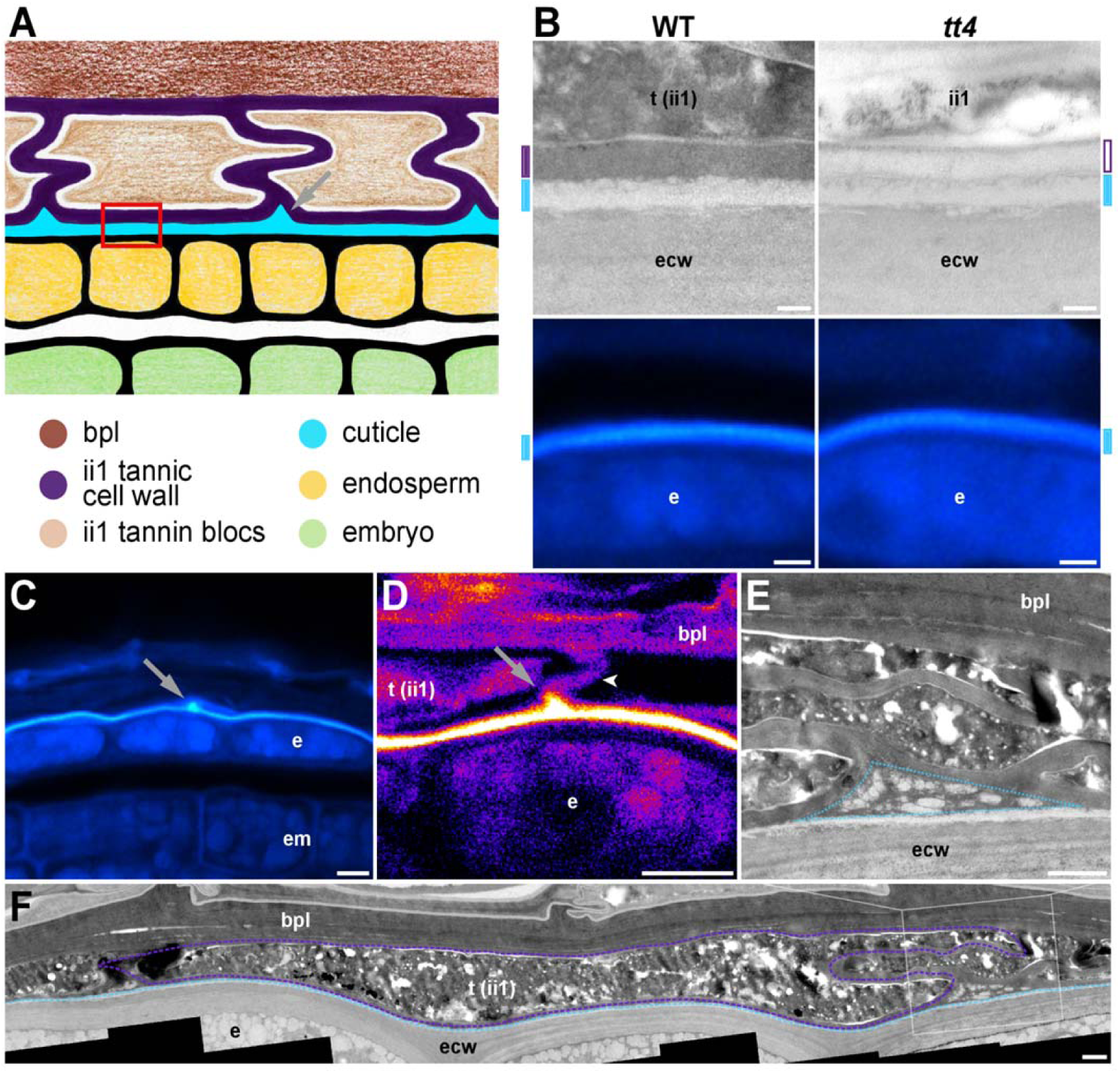
ii1 cell walls are tannified and form a continuous reticulated tannic apoplastic barrier tightly bound to the endosperm-associated cuticle. (A) Schematic drawing illustrating the organization of the endosperm outer surface area. The drawing is not made to scale in order to better visualize the positions of the different structures of interest. The red-boxed region indicates the surface of the endosperm where the cuticle and inner tannified cell wall are tightly associated. The grey arrow indicates the location of cuticular accumulation. (B) TEM micrographs of the endosperm outer surface region (top row) and paraffin sections stained with Auramine O (bottom row) in WT and *tt4* mutants, corresponding to the red-boxed region shown in A. Cyan marks indicate the position of the cuticle and purple marks indicate the position of the ii1 inner periclinal cell wall (filled in purple and in white when it is electron-dense and electron-lucent, respectively). (C) Auramine O staining of a WT seed paraffin section; the arrow indicates a local signal accumulation corresponding to the region indicated by the grey arrow in A. (D) Confocal micrograph of a WT seed paraffin section stained with Auramine O. The arrow indicates a local signal accumulation corresponding to the region indicated by the grey arrow in A; the arrowhead indicates an anticlinal cell wall between two adjacent ii1 cells. (E) TEM micrograph of a junction between ii1 cells in a WT seed (enlargement of the white-boxed region in F). The dashed blue triangle indicates the region where droplet-like electron-lucent material accumulates, between neighboring ii1 cells and the cuticle. (F) TEM micrograph presenting a global view of the WT seed inner integuments. The purple dashed line indicates the cell wall of one ii1 cell (electron-dense material) and the light blue dashed line indicates the cuticle (electron-lucent material). The original picture is in Supplemental Figure S5A. Scale bars: B, 200 nm and 2 μm, top and bottom rows, respectively; C and D, 5 μm; E and F, 1 μm. Abbreviations: bpl, brown pigment layer; e, endosperm; ecw, endosperm cell wall; em, embryo; ii1, inner integument 1; t, tannins.

To evaluate this model, we first assessed whether the electron-dense reticulated structure corresponds indeed to tannified cell walls. Interestingly, the linear structure revealed by DMACA at the surface of the endosperm periodically protruded outwardly, forming a reticulated extended structure in a manner most similar to that observed with the electron-dense layer in TEM (arrowheads in Fig. 1C, left; Supplemental Fig. S3A). Furthermore, DMACA did not reveal a reticulated structure in *tt4* mutant seed sections (Fig. 1C, right panels). In addition, the electron-dense reticulated structure was no longer electron-dense in *tt4* mutants (Fig. 2B, top row). These observations strongly indicate that reticulated structure revealed by DMACA corresponds to the electron-dense structure observed in TEM.

Calcofluor white (CW) is a fluorescent dye used for staining plant cell walls due to its specific binding to cellulose (Hughes and McCully, 1975). Semi-thin WT seed sections stained with CW revealed the presence of linear plant cell wall material periodically protruding outwardly from the surface of the endosperm, consistent with the notion that it corresponds to ii1 cell wall material (arrowheads in Supplemental Fig. S4). Furthermore, the CW signal formed a reticulated structure reminiscent of the one observed with DMACA and TEM (Supplemental Fig. S4). Altogether, these observations show that the electron-dense reticulated structure indeed corresponds to tannified cell walls.

We next assessed whether the electron-lucent layer indeed corresponds to the endosperm-associated cuticle. We observed that the electron-lucent layer had the same appearance in both WT and *tt4* mutants, consistent with previous reports (Fig. 2B, top row) (Loubéry et al., 2018). Similarly Auramine O, a fluorescent cuticle marker (Lequeu et al., 2003; Szczuka and Szczuka, 2003; Buda et al., 2009), revealed the presence of a linear signal that covers the outer surface of the endosperm; furthermore, it had the same appearance in both WT and *tt4* seeds (Fig. 2B, bottom row). Thus, unlike the electron-dense/DMACA structure, the appearance of the electron-lucent/Auramine O layer is not perturbed by the absence of tannins, indicating that they correspond to distinct structures. Furthermore, these results strongly support the notion that the electron-lucent layer and the Auramine O signal represent the same structure, namely the endosperm-associated cuticle.

To further ascertain this claim, we took advantage of an unexpected observation. Indeed, we observed that the Auramine O linear signal was occasionally punctuated by patches protruding outwardly (Fig. 2C). Confocal micrographs suggested that these patches are present at the meeting location of two tannic cell walls from adjacent ii1 cells (Fig. 2D). If the electron-lucent material indeed corresponds to the Auramine O structure, then the existence of Auramine O patches predicts that corresponding electron-lucent depositions should be visible in TEM and that they should be located at the meeting point of adjacent ii1 cells.

Accordingly, TEM images confirmed that at the meeting point of two adjacent ii1 cells, electron-lucent material accumulated on the outer side of the cuticle (Fig. 2, E and F). Indeed, after protruding out from the surface of the endosperm and before fusing, two tannic cell walls from adjacent ii1 cells form two edges of a triangle whose third edge is the electron-lucent layer that remains associated with the surface of the endosperm (Fig. 2, E and F). Electron-lucent material accumulated in the interior of the triangle, taking the form of an assembly of droplet-like structures (Fig. 2E). We noticed that the number and surface of these droplet-like structures varied considerably among the various triangular locations, and they were sometimes absent (Supplemental Fig. S5B). This is consistent with the fact that the patches of signal revealed by Auramine O were also not systematically visible along the outer surface of the endosperm. These observations provide further correlative evidence that the electron-lucent layer indeed corresponds to the endosperm-associated cuticle.

Altogether, we conclude that the outer endosperm surface is covered by a two-layered structure consisting of a *bona fide* cuticle and a tannic cell wall, both derived from ii1 cells. ii1 cell walls are tannified and further protrude outwardly, forming a continuous tannic reticulated apoplastic barrier surrounding the seed’s living tissues. They support the notion that the tannic cell wall is a previously unrecognized building block of the Arabidopsis mature seed coat.

### The ii1 inner periclinal tannic cell wall remains attached to the endosperm after seed coat rupture

We previously showed that the outer surface of the endosperm is more permeable to toluidine blue in both *bodyguard* mutants, deficient in cuticle formation, and *tt4* mutants (De Giorgi et al., 2015; Loubéry et al., 2018). Together with the model proposed above, these observations would therefore support the notion that both the cuticle and the tannic cell wall contribute to the biophysical properties of the outer surface of the endosperm. However, to ascertain this claim we sought to verify that both maternal structures indeed remain present at the outer surface of the endosperm upon seed coat rupture, which was not previously investigated.

In unstained WT seeds undergoing seed coat rupture, the exposed outer endosperm surface was covered by an unanticipated and striking pattern of brown-edged polygons, which was not previously reported (Fig. 3A). When WT seeds were stained with DMACA the polygonal pattern was even more striking, as DMACA strongly colored the polygon edges (Fig. 3B, left panel). Furthermore, the interior of the polygons was also lightly stained by DMACA and this coloration was not visible in *tt4* mutants (Fig. 3B, right panels). These observations provided a first indication that the surface of the endosperm remains entirely covered by ii1 inner periclinal tannic cell walls during seed coat rupture, notably as manifested by the tannin-specific DMACA coloration of the polygon interiors.

**Figure 3.**
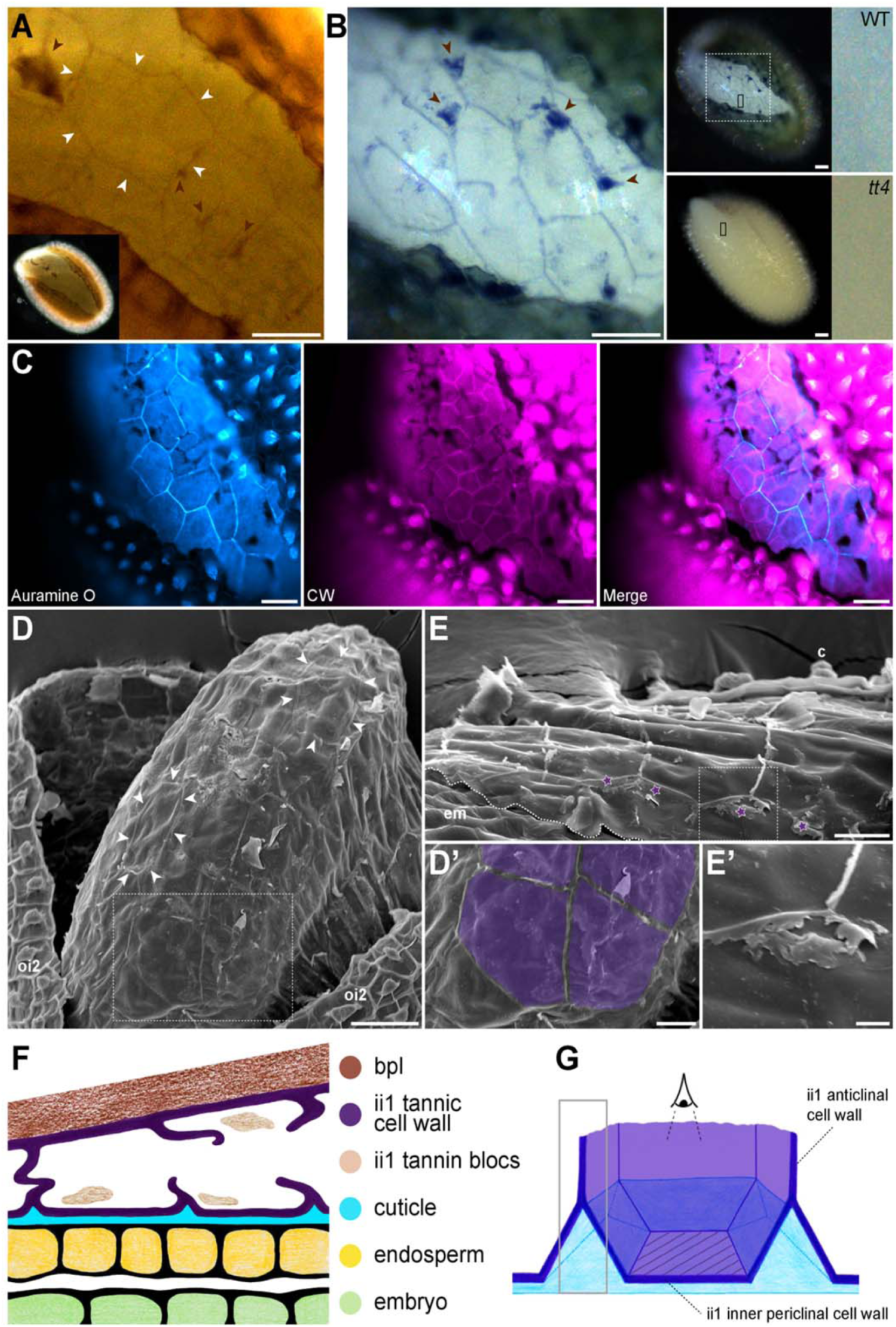
The ii1 inner periclinal tannic cell wall remains attached to the endosperm after seed coat rupture. (A) Unstained seed coat-ruptured WT seed showing a polygonal pattern present at the outer surface of the endosperm. White arrowheads delineate a particular polygonal contour. (B) DMACA staining of seed coat-ruptured WT and *tt4* seeds. The left panel is an enlargement of the dashed-boxed area shown in the WT seed picture of the middle panel. The right panels are enlargements of the small black boxes shown in the middle panel. In A and B, brown arrowheads indicate remnants of ii1 tannin blocks (see Discussion). (C) Seed coat-ruptured WT seed co-stained with Auramine O and CW. (D and E) SEM micrographs of seed coat-ruptured (D) and endosperm-ruptured (E) WT seeds. White arrowheads indicate the visible remnants of ii1 cell walls. (D’) Enlargement of the boxed region in D, where ii1 inner periclinal wall surfaces were colorized in purple. (E’) Enlargement of the boxed region in E, where remnants of ruptured ii1 anticlinal cell walls are visible (indicated by purple stars in E). Note that SEM was performed on fresh (non-fixed) material; because of the microscope vacuum, ruptured seed coats would close themselves, hiding zones of interest (see Supplemental Figure S8B for an example); to remedy this, we had to manually increase the aperture of seed coat-ruptured seeds, thus uncovering a wider space than that corresponding to a normal seed coat-ruptured seed. (F) Schematic drawing of seed coat organization during seed coat rupture (outer integuments are not represented). (G) Schematic drawing in 3D of one ii1 cell during seed coat rupture. The grey-boxed region corresponds to the signal given by DMACA (in B) and Auramine O (in C) along broken anticlinal ii1 walls. The hatched area indicates the ii1 inner periclinal cell wall surface, which becomes the first barrier facing the environment during seed coat rupture. Drawings (F and G) were not made to scale in order to better visualize the structures of interest. Scale bars: A-D, 50 μm; D’ and E, 20 μm; E’, 5 μm. Abbreviations: c, columella; em, embryo; ii1, inner integument 1; oi2, outer integument 2.

The presence of a pattern of brown-edged polygons at the outer surface of endosperm cells, further exacerbated by DMACA, strongly suggested that anticlinal ii1 tannic cell wall material remains attached to the ii1 inner periclinal tannic cell wall. Hence, the polygonal pattern would result from additional ii1 anticlinal cell wall material, which in turn would generate an imprint of the ii1 cell contours on the outer surface of the endosperm (Fig. 3, A and B). Incidentally, this suggestion would imply that the anticlinal ii1 tannic cell walls are the rupture points during seed coat rupture, which was not previously reported. We sought to further evaluate these claims by monitoring cuticular material at the surface of the endosperm. Indeed, we saw above that patches of cuticular accumulation are found on the outer side of the cuticle where two adjacent ii1 cells meet (Fig. 2, C-E). Therefore, the model predicts that this cuticular accumulation should also generate a polygonal pattern at the outer surface of endosperm cells corresponding to an imprint of ii1 cells.

Seeds that had undergone seed coat rupture were stained with the cuticle dye Auramine O. As anticipated by the model, this also revealed a striking polygonal pattern at the outer surface of the endosperm similar to that seen with DMACA (Fig. 3C, left panel). Demonstrating that the Auramine O polygonal pattern matches a polygonal pattern generated by ii1 cell wall would require co-staining whole mount seed coat-ruptured seeds with Auramine O and CW to show that the polygonal edges coincide. However, as illustrated in Supplemental Figure S4, ii1 cell walls are very weakly stained with CW compared to endosperm cell walls; in addition, the very small distance between the endosperm outer cell wall and the ii1 inner periclinal cell wall is below the resolution of the light microscope. As a result, it is expected that in whole-mount stainings, the signal given by CW on ii1 inner periclinal cell walls will be hidden by the much stronger signal given by the endosperm outer cell wall. Indeed, CW enabled visualizing endosperm cell walls and, deeper within the seed, embryo cell walls, but it did not reveal ii1 cell walls (Fig. 3C, middle panel; Supplemental Fig. S6).

We therefore opted for an indirect approach. The polygonal pattern revealed by Auramine O at the outer surface of the endosperm corresponds to either the anticlinal endosperm cell walls or the anticlinal ii1 cell walls. Thus, if the polygonal pattern revealed by Auramine O corresponded to endosperm anticlinal walls, it would co-localize with the signal given by these walls upon CW staining. Strikingly, the Auramine O polygonal pattern was seen in close vicinity to endosperm cell walls and slightly above them (Fig. 3C). Furthermore, their respective patterns did not match, demonstrating that the Auramine O signal does not delineate endosperm cells (Fig. 3C, right panel). This is further evidenced by the fact that endosperm cells and ii1 cells have a substantial size difference: ii1 cells are significantly larger than endosperm cells (Figs. 1C and 2F; Supplemental Figs. S1-S4). This size difference was indeed observed between the contours delineated by DMACA and Auramine O, and those of endosperm cells revealed by CW (Fig. 3, B and C). Hence, we conclude that the polygonal pattern revealed by Auramine O corresponds to an imprint of ii1 cells at the outer surface of the endosperm.

Altogether, these data are strongly consistent with the notion that the endosperm-associated cuticle and the inner periclinal tannic ii1 cell walls remain attached to the endosperm surface upon seed coat rupture.

To further corroborate this claim, we used scanning electron microscopy (SEM) on seed coat-ruptured seeds. Again, a polygonal pattern could be recognized at the surface of the endosperm (Fig. 3, D and D’; Supplemental Fig. S7). The polygons matched in size and shape those described above without dye or with DMACA and Auramine O (Fig. 3, A-C). Furthermore, in some regions of the outer endosperm surface, underlying endosperm cells collapsed partially, likely as a result of the vacuum of the SEM: this generated a hollow pattern that revealed the contour of endosperm cells. The endosperm cell contours were significantly smaller than the polygons observed at their surface (Supplemental Fig. S8A). These observations are consistent with the model described above, namely that ii1 anticlinal cell walls remain attached at the outer surface of the endosperm leaving an imprint of ii1 cell contours.

This conclusion is further supported by the following observation: close inspection revealed that the polygonal edges observed in SEM had sheet-like appearance and quality (Fig. 3, E and E’). The extension of this sheet-like material varied along the polygonal contours. These observations support the notion that the sheet-like material corresponds to ruptured ii1 anticlinal cell walls, which leaves, as a result, portions of cell walls of different sizes resting on the ii1 inner periclinal cell walls that cover the endosperm-associated cuticle (Fig. 3F).

Altogether, these observations corroborate the notion that during seed coat rupture, the endosperm-associated cuticle and the ii1 inner periclinal tannic cell wall remain attached to the outer surface of the endosperm after ii1 anticlinal cell wall rupture (Fig. 3F). The juxtaposed cuticle and ii1 inner periclinal tannic cell wall two-layer constitutes the first physical barrier separating living tissues from the outer environment (Fig. 3, F and G).

Together with the toluidine blue permeability experiments reported by De Giorgi et al. and Loubéry et al. (De Giorgi et al., 2015; Loubéry et al., 2018), these observations show that both the cuticle and the tannic cell wall of this two-layer structure contribute to the biophysical properties of the outer surface of the endosperm.

The identification of a tannic cell wall directly involved in the permeability properties of the Arabidopsis mature seed coat prompted us to further document its occurrence in other parts of the seed coat.

### The brown pigment layer is mostly composed of tannic cell walls

During seed maturation, differentiating ii1’ and ii2 layers form the brown pigment layer (bpl). TEM observations indicated that the bpl is essentially composed of cell walls (Fig. 2, E and F). Indeed, no (or very little) cytoplasmic remains were visible in the ii1’ and ii2 layers (Fig. 2, E and F; Supplemental Fig. S2), and the bpl appeared to be formed by the close apposition of ii1’ and ii2 cell walls with the outer ii1 and the inner oi1 cell walls. This conclusion is further supported by CW staining, showing extensive labeling of cellulose throughout the bpl (Supplemental Fig. S4).

Furthermore, the TEM observations in mature seeds reported here show that ii1 cell walls and the bpl have a similar electron density so that their delimitation is sometimes difficult to discern, suggesting that the bpl might also be made of cell walls impregnated by tannins (Fig. 2, E and F; Supplemental Fig. S2). Consistent with this view, in *tt4* seeds the bpl had an electron-lucent appearance in TEM similar to that of ii1 cell walls (Supplemental Fig. S9). In addition, in unstained *tt4* histological sections, the bpl lost its brown coloration (Supplemental Fig. S1, B and C).

The hypothesis of the bpl as a tannin-rich structure is further and strongly supported by the clear signal given by the bpl upon staining with the tannin-specific dye DMACA (Fig. 1C; Supplemental Fig. S3). Indeed, DMACA stainings of semi-thin sections showed a strong signal in the entire bpl layer, similar to that observed in the ii1 layer; furthermore, this signal was absent in *tt4* mutant seeds (Fig. 1C; Supplemental Fig. S3B). These observations show that tannins are not only present in the ii1 layer but also in the bpl.

In WT seed sections (unstained, or upon staining with toluidine blue or methylene blue), the bpl did not have exactly the same color as that of the interior of ii1 cells and ii1 cell walls (Fig. 1, A and B; Supplemental Fig. S1, A-C); however all these structures were recognized by DMACA, attesting that tannins are part of their composition. The variability in color may be due to additional components present in the bpl, such as other types of flavonoids (e.g. flavonols).

Thus, our observations show that tannic cell walls are major building blocks of the bpl. By extension, and taking into account the juxtaposition of ii1 tannic cell walls and the bpl, they shed a new light on tannic cell walls, which appear to account for a substantial part of the mature Arabidopsis seed coat.

## DISCUSSION

*tt* mutant seeds have low viability and dormancy due to a defective seed coat. We sought to understand the origin of the intriguing permeability defects observed in the endosperm-associated cuticle of *tt* mutants (*tt4, tt5, ban, tt12* and *tt15*) reported by Loubéry et al., 2018. Indeed, *tt* mutants are deficient not in cuticular components synthesis, but rather in flavonoid synthesis. This led us to identify here a tannic cell wall tightly associated with the endosperm-associated cuticle; furthermore this component remains attached to the cuticle at the surface of the endosperm during seed coat rupture where it contributes, together with the cuticle, to the permeability properties of the endosperm. Thus, we interpret the permeability defects of the endosperm in *transparent testa* mutants reported by Loubéry et al., 2018 as the result of the absence of tannins in the cell wall that is present, together with a cuticle, at the surface of the endosperm. Furthermore, we show that tannic cell walls are the main constituents of the bpl, therefore identifying them as a major structural component of the mature seed coat. In summary, we report the identification of tannic cell walls as previously unrecognized seed coat structures that form a continuous barrier in the seed coat and contribute to the biophysical properties of the seed.

### Reinterpretation of the two-layered structure observed in TEM at the outer surface of endosperm cells

In a previous report, the two-layered structure observed in TEM at the outer surface of endosperm cells, consisting of an inner electron-lucent layer and an outer electron-dense layer was viewed as a cuticle made of two different compositions (De Giorgi et al., 2015; Loubéry et al., 2018).

In the scientific literature, cuticles are predominantly described as electron-dense layers, according to their appearance in TEM, whether in seeds, stem or leaves (Beeckman et al., 2000; Andème Ondzighi et al., 2008; Yang et al., 2008; DeBolt et al., 2009; Voisin et al., 2009; Bowman, 2011; Shumborski et al., 2016; Moussu et al., 2017). However, previous reports have described amorphous electron-dense procuticles that are converted to a simple electron-lucent amorphous cuticle (Jeffree, 2007). Thus, the electron-dense character of a layer observed in TEM is not an absolute criterion to define a cuticle and several cuticles have been described as partially or entirely electron-lucent (Jeffree, 2007; Budke et al., 2011; Guzmán et al., 2014; Bourgault et al., 2020). Interestingly, examining throughout development the two-layer structure that borders the endosperm brings a picture that is rather consistent with a model of a cuticle progressively losing its electron-dense character. Indeed, the endosperm-associated two-layered structure is of maternal origin, deriving from ii1 cells and transferred to the endosperm between the torpedo and walking-stick stage (Loubéry et al., 2018). During earlier stages of seed development (globular and heart stages) the structure is present at the surface of ii1 cells and is made of two layers but with an inverted electron density relative to that found in mature seeds. Indeed, at early stages the cuticle is electron-dense whereas the ii1 cell wall is rather electron-lucent (Loubéry et al., 2018). From the mature embryo stage onwards the cuticle is no longer electron dense; in contrast, the ii1 cell wall becomes electron-dense only at late seed maturation stages (see below).

Thus, we conclude that, in mature seeds, of the two layers at the outer surface of the endosperm only the inner layer is made of cuticular material, the outer layer corresponding to tannins-impregnated ii1 cell wall as further discussed hereafter.

### Identification of a new structure in the seed: the tannic cell wall

All the ovular integument layers of the *Arabidopsis* seed coat differentiate and ultimately die in the course of seed development according to distinct temporal developmental programs (Ingram, 2017). ii1’ and ii2 cell layers present the first signs of terminal differentiation (nuclei degradation, vacuolar disorganization and cell shrinkage) during the heart and torpedo stages of embryo development (Nakaune et al., 2005). Their death leads to the formation of a condensed structure in the seed coat referred to as the brown pigment layer (bpl).

It has been reported that the *Arabidopsis* mature seed coat contains two types of flavonoids: flavonols in the outer integument 1 (oi1) layer (Pourcel et al., 2005; Routaboul et al., 2006) and tannins mainly in ii1 cells, and also in a few cells in the chalazal and micropylar areas (Debeaujon et al., 2003, 2007). Here we show that tannins are additionally present in ii1 cell walls. We also show that the bpl is, remarkably, constituted of cell walls, themselves also impregnated with tannins.

Tannin precursors are biosynthesized in vesicles arising from the endoplasmic reticulum before their incorporation into vacuoles (Stafford, 1988; Kitamura et al., 2004). The *BANYULS* gene, which encodes an enzyme specifically involved in tannin biosynthesis, is majoritively expressed in the ii1 cell layer during early stages of seed development (Devic et al., 1999; Debeaujon et al., 2003). Additionally, the expression of other genes involved in tannin biosynthesis, transport and compartmentalization was also detected in ii1 cells, but not in ii1’ or ii2 (Debeaujon et al., 2001; Pourcel et al., 2005; Marinova et al., 2007). Based on these data, we propose that after being synthesized in ii1 cells, tannins, or their precursors, migrate from their cytosolic vacuole to the apoplast, impregnating progressively ii1 as well as ii1’ and ii2 cell walls that form the bpl layer. These events would occur following ii1 cell death, which takes place at late stages of seed development and induces the burst of vacuoles and the liberation of their tannic content.

Eventually, in the mature seed, ii1 tannic cell walls together with the bpl tannic cell walls define a large and continuous electron-dense tannin-rich region in the seed coat (Supplemental Figs. S2 and S5). Two electron-lucent and previously characterized impermeable barriers border this structure: on the inner side, the endosperm-associated cuticle and on the outer side, a suberized cell wall (Yadav et al., 2014; De Giorgi et al., 2015; Gou et al., 2017; Loubéry et al., 2018). The presence of these two barriers may explain why tannins synthesized in ii1 cells diffuse to the bpl during seed development, but not further. Furthermore, tannins shield living tissues from external damage, but they in turn could interfere with vital functions: hence tannin containment within the seed coat may be crucial to avoid exposing living tissues to these potentially harmful compounds.

### Previous evidence for tannic cell walls

Several reports have described the presence of flavonoids in cell walls (Dai et al., 1996; Dünisch et al., 2010; Ermeydan et al., 2012) or in cuticles (Domínguez et al., 2009; España et al., 2014). Concerning tannins, in most reported cases, tannic compounds are sequestered within cells, in the vacuolar compartment (Abrahams et al., 2003; Lepiniec et al., 2006; Debeaujon et al., 2007; Marinova et al., 2007; Petrussa et al., 2013; Hammouda et al., 2014; Zhang et al., 2019). However, the presence of tannins in cell walls has been proposed in roots of various plant species (Evert, 2006). Concerning the presence of tannic cell walls in seeds, Kitamura et al. observed a weak tannin signal revealed by vanillin outlining ii1 cells in mature seeds; however, the resolution of the micrographs in the study did not allow to assess whether tannins were situated inside ii1 cells or in their walls (Kitamura et al., 2004).

### Proposed functions of tannins in the seed coat

In Arabidopsis seeds the absence of flavonoids correlates with higher seed permeability to water and oxygen; in turn, this lowers seed dormancy and viability (Debeaujon et al., 2000; Chahtane et al., 2017). Tannins can act as antioxidants and may thus limit oxygen diffusion in the seed coat (Harborne and Williams, 2000; Pietta, 2000; Debeaujon et al., 2007; Pourcel et al., 2007). Furthermore, some tannins are known to be indeed impermeable to water (Debeaujon et al., 2007). However it was unclear so far how tannins, which appear as discontinuous blocks inside ii1 cells, could act as a permeability barrier. The characterization here of tannic cell walls sheds light on how flavonoids could act as a permeability barrier: indeed, they form a continuous barrier surrounding the seed’s living structures, thus shielding them from the outer environment. Furthermore, the permeability experiments reported by Loubéry et al., 2018, showing that *tt* mutant endosperm is more permeable to toluidine blue in seed coat-ruptured seeds, together with the evidence presented here that the tannic cell wall remains associated with the endosperm upon seed coat rupture, provides direct evidence that the tannic cell wall indeed renders the seed coat less permeable to outer compounds.

In seeds deprived of tannins, such as *tt4* mutant seeds, the ii1 cell layer is considerably more collapsed than in WT seeds (Fig. 1, B and C), which is consistent with the thinner and weaker seed coats of *tt4* mutant seeds (Debeaujon et al., 2000). Interestingly, it is known that tannins interact with cell wall components, by making cross-links with proteins and polysaccharides (Marles et al., 2003; Debeaujon et al., 2007), thus strengthening the whole structure. The presence of tannins in ii1 cell walls could then increase the mechanical stability of these cells, and globally enhance the hardiness of the seed coat. Furthermore, it has been demonstrated that artificial incorporation of flavonoids in wood cell walls improves wood stability upon changes in temperature and humidity; it also enhances its durability by reducing the moisture content, hence decreasing the risk of degradation by fungi (Ermeydan et al., 2012). This suggests that the presence of tannins in ii1 cell walls and the bpl may indeed improve Arabidopsis seed solidity and durability.

The list of proposed or demonstrated flavonoids functions is extensive: flavonoids have been shown in particular to act as signaling molecules between plants and other organisms, delivering messages of attraction, repulsion or inhibition. Arabidopsis plant roots are known to maintain relations with microbial communities, which can positively affect their health and development (Bergelson et al., 2019; Cheng et al., 2019b). As it is established that flavonoids can act as signaling molecules with soil organisms, we can speculate that tannins from the seed coat could prime the development of these symbiotic relations for the upcoming roots. Interestingly, we showed that seed coat rupture occurs at the level of anticlinal ii1 cell walls, which is expected to release the tannins they contain. Consistent with this notion, patches stained by DMACA are occasionally detectable on the endosperm surface upon seed coat rupture (brown arrowheads in Fig. 3B and in Supplemental Fig. S10), but in most cases they do not appear to remain attached to the surface of the endosperm, suggesting that they are released in the seed environment. Thus, it is tempting to speculate that if tannins were to have a role in the establishment of mutualistic interactions with microorganisms, the mechanics of seed coat rupture would directly time the release of tannins with the moment when they are needed. In addition, upon imbibition, seeds can suffer sudden abiotic stresses, such as dehydration, an osmotic disequilibrium or variations of temperature, which leads to a germination arrest until conditions are favorable again (Lopez-Molina et al., 2001, 2002). Germination arrest could then take place in seeds that already ruptured their seed coat. During this intermediate developmental stage the two-layer made of a cuticle and a tannic cell wall, together with the release of the ii1 cell tannin contents, may maintain a certain level of impermeability to water and oxygen, as well as some resistance to pathogens and herbivores.

Another widespread role for flavonoids (including tannins) that has been described in various species is in the defense against herbivores or pathogenic bacteria and fungi. Tannins toxicity is due to their complexation with proteins or substrates, alteration of membranes or complexation with metal ions, which causes severe limitations for pathogen health and growth (Scalbert, 1991; Chung et al., 1998; Shirley, 1998; Dixon et al., 2005). Seeds are known to be the target of deleterious fungi and bacteria: indeed, seeds infected with fungi or bacteria deteriorate faster and have a lower viability than uninfected seeds (Mohamed-Yasseen et al., 1994). One putative role of tannic walls in the seed could be to shield the seed against soil microorganisms so as to protect the embryo from pathogenic attacks. Concerning bigger predators, it has been shown that some animals avoid seeds with a high tannin content, likely due to the astringency of tannins, and in consequence either avoid consuming seeds with a high tannin content, or else reject the seeds after fruit consumption (Harborne and Williams, 2000; Zhang et al., 2013; Ancillotto et al., 2015). This might be beneficial for the plant as well as for the consumer. Indeed, in both cases the seed is not destroyed by the animal and its dispersion is facilitated.

### Development and presence of different protective layers in the seed coat

In the Arabidopsis seed coat, several layers participate in the protection of the embryo and the endosperm. In order to face various sources of stress, these layers have different compositions: a cuticle along the endosperm surface; tannins inside ii1 cells and in their walls, as well as in the bpl; suberin and flavonols in oi1 cells (Pourcel et al., 2005; Yadav et al., 2014; Gou et al., 2017). We showed that from seed coat rupture onwards, the endosperm is covered by a newly identified two-layered maternal structure made of a cuticle and a tannic cell wall, deriving from ii1 cells, and conferring a supplemental level of defense and protection. Both components appear to have similar functions, in particular for the control of water and oxygen passage, ensuring a high level of protection in this regard. Additionally, tannins released by ii1 cells upon seed coat rupture may bring additional putative functions regarding the interactions of Arabidopsis seeds with pathogenic or beneficial organisms, which remains to be investigated.

## MATERIAL AND METHODS

### Plant material and growth conditions

Arabidopsis (*Arabidopsis thaliana*) Columbia (Col-0) seeds were used as wild-type material together with the *tt4-5* mutant in the Col-0 background (NASC stock number N66121, contributed by F. Zhang, Y. Qi and D. Voytas). For each experiment presented, the seed material used (i.e. the wild-type seed material and the mutant seed material) was harvested on the same day from plants grown under identical environmental conditions. Dry siliques were obtained around 8 weeks after planting.

### Semi-thin sections

8 h-imbibed seeds were delicately punctured with a fine needle and fixed overnight at 4°C in 2.5% (v/v) glutaraldehyde and 0.01% (v/v) Tween-20 in phosphate buffer (pH 7.2) after vacuum infiltration. Samples were then washed with water, embedded in pellets of 1.5% (w/v) agarose, dehydrated in a graded ethanol series, and embedded in Epon 812. Finally, 1 μm semi-thin sections were cut using a UCT microtome (Leica) and placed on SuperFrost slides. For DMACA stainings, seed sections were incubated in the dark during 72 h at room temperature in a solution of 0.3% (w/v) DMACA (Sigma-Aldrich) in a mixture of methanol and 6 M HCl (1:1, v/v). To prepare this solution the DMACA powder was first solubilized in methanol and agitated for 10 minutes, then H_2_O and HCl were added to the mixture. The slices were finally rinsed several times with 50% ethanol and mounted in water. For toluidine blue staining, fixed seed sections were incubated in a solution containing 0.5% (w/v) toluidine blue in 0.1 M phosphate buffer (pH 6.8) during 5 min at 70°C on a hot plate, rinsed with H_2_O and mounted in Eukitt. For methylene blue staining, a stock solution was prepared containing 0.15% (w/v) methylene blue in 10% glycerol, 10% methanol, 30% distilled water and 50% phosphate buffer (pH 6.9); the solution was then diluted 20x in distilled water and fixed seed sections were incubated during 5 min at 70°C on a hot plate, rinsed with H_2_O and mounted in Eukitt. For Calcofluor White stainings, sections were first incubated for 2 min in 0.01% (w/v) Calcofluor White in water, then washed with PBS (pH 6.8) and mounted in PBS (pH 6.8) with glycerol (1:1, v/v). In Figure 1A (center bottom) and Supplemental Figure S1, B and C, non-stained slices were simply mounted in Eukitt.

### Paraffin sections

8 h-imbibed seeds were delicately punctured with a fine needle and fixed overnight at 4°C in phosphate buffer (pH 7.2) with 4% (v/v) formaldehyde and 0.25% (v/v) glutaraldehyde, after vacuum infiltration. Samples were then embedded in pellets of 1.5% (w/v) agarose, dehydrated in a graded ethanol series, cleared in Neoclear, and embedded in paraffin. 8 μm-thick sections were cut with a Cut 4050 microtome (MicroTec), placed on SuperFrost slides (Roth), deparaffinized with Neoclear, and rehydrated with water. For Auramine O staining, sections were incubated for 5 min in 0.001% (w/v) Auramine O in water, then washed with PBS (pH 6.8) and mounted in PBS (pH 6.8) with glycerol (1:1, v/v).

### Whole-mount stainings

Fresh seeds were plated on MS medium supplemented with 10 μM ABA during 36 h (until seed coat rupture occurred). For double Calcofluor White/Auramine O stainings, seeds were first incubated for 20 min in 0.01% (w/v) Calcofluor White in water, then washed with water and incubated for 30 min in 0.005% (w/v) Auramine O in water; finally, they were washed with water and mounted in water between slide and coverslip. For DMACA stainings, seeds were incubated during 1 h in a DMACA solution (see above), rinsed several times with 50% ethanol and mounted in water between slide and coverslip. In Figure 3A, non-stained seeds were simply mounted in water between slide and coverslip.

### Light microscopy

Samples were examined with an Eclipse 80i widefield microscope (Nikon) equipped with a 40x Plan Fluor NA 0.75 air lens and a Digital Sight DS-Fi1 color CCD camera (Nikon) or a DS-Fi3 CMOS color camera (Nikon). Fluorescence excitation was done with an Intensilight C-HGFI mercury vapor lamp (Nikon); Calcofluor White was examined using a 4′,6-diamino-phenylindole filter set (excitation, 352–402 nm; emission, 417–477 nm), Auramine O was examined using a CFP filter set (excitation, 426–450 nm; emission, 467–499 nm) and for Figure 1A, autofluorescence was examined using a GFP long-pass filter (excitation, 460-500 nm; emission > 510 nm). For Figure 2D, Auramine O was examined using an SP5 confocal microscope (Leica) equipped with a 63x PlanApo NA 1.4 oil lens, an argon laser with excitation at 458 nm, a HyD detector with emission collection between 483 and 513 nm, and using a pixel size of 100 nm; a maximal projection of a few z planes was performed and displayed. Whole-mount samples in Figure 3B and Supplemental Figure S10 were examined with an Axio Zoom.V16 stereomicroscope (Zeiss) equipped with a 1x NA 0.25 objective and an Axiocam 512 CMOS camera, under episcopic illumination.

### TEM

TEM was performed as described previously (Loubéry et al., 2018). Briefly, 8 h-imbibed seeds were delicately punctured with a fine needle and fixed overnight at 4°C in 2.5% (v/v) glutaraldehyde and 0.01% (v/v) Tween-20 in 100 mM sodium cacodylate (pH 7) after vacuum infiltration. After a primary postfixation in 1.5% (v/v) osmium tetroxide for 2 h at 4°C and a secondary postfixation in 1% (w/v) uranyl acetate for 1 h at 4°C, seeds were embedded in pellets of 1.5% (w/v) agarose, dehydrated in a graded ethanol series, and embedded in Epon 812. Then, 85 nm ultra-thin sections were cut using a UCT microtome (Leica), stained with 2.5% (w/v) uranyl acetate and Reynolds lead citrate, and finally observed with a Tecnai G2 Sphera (FEI) at 120 kV, equipped with a high-resolution digital camera.

### SEM

Fresh seeds were plated on MS medium supplemented with 10 μM ABA during 36 h (until seed coat rupture occurred). Seeds were delicately transferred onto SEM holders and placed on double-sided carbon tape (Electron Microscopy Sciences), and afterwards treated with a gold sputter coater (JFC-1200, JEOL). Imaging was performed with a JSM-6510LV (JEOL) scanning electron microscope in high vacuum mode with a spot size of 40, an acceleration of 15 kV and a working distance of 12 mm.

### Image treatment and analysis

Images were treated and analyzed using the software Fiji (Schindelin et al., 2012). For both histology and TEM, tiling and stitching were used to obtain high resolution large fields of view. Tiling was performed manually, and stitching was done using the Fiji MosaicJ plugin (Thévenaz and Unser, 2007). The left panel in Figure 1A was modified using Adobe Photoshop Lightroom CC version 2015.12 and Adobe Photoshop CC 2017. In Figure 3C and Supplemental Figure S3A, Calcofluor White stainings were colorized with the Fiji Magenta and Grays look-up tables, respectively; in Figure 2D, Auramine O staining was colorized with the Fiji Fire look-up table, to provide a larger dynamic range and a higher quality of visualization. All figures were mounted using Adobe Illustrator CC 22.0.0.

## ACKNOWLEDGMENT

This work was supported by Swiss National Science Foundation grants to LLM (grant nos. 31003A-152660/1 and 31003A-179472/1) and the State of Geneva, Switzerland.

---

**Table 1.**
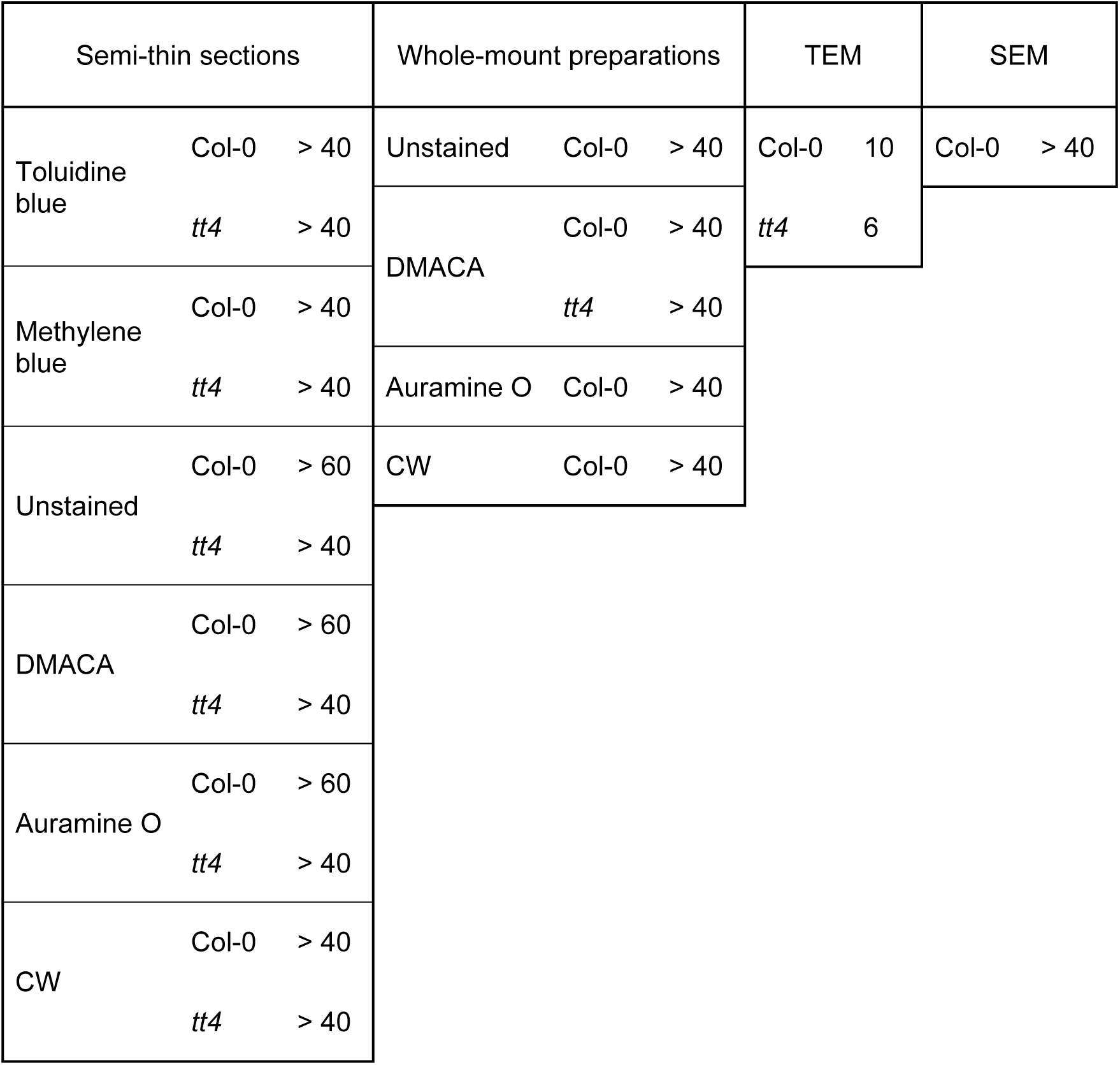
Biological sample sizes. The table presents the seed numbers analyzed for each set of experiments and genotypes presented in the article. Each experiment was performed at least in triplicate.

| Semi-thin sections |  |  | Whole-mount preparations |  |  | TEM |  | SEM |  |
| --- | --- | --- | --- | --- | --- | --- | --- | --- | --- |
| Toluidine blue | Col-0 | > 40 | Unstained | Col-0 | > 40 | Col-0 | 10 | Col-0 | > 40 |
|  | *tt4* | > 40 | DMACA | Col-0 | > 40 | *tt4* | 6 |  |  |
| Methylene blue | Col-0 | > 40 |  | *tt4* | > 40 |  |  |  |  |
|  | *tt4* | > 40 | Auramine O | Col-0 | > 40 |  |  |  |  |
| Unstained | Col-0 | > 60 | CW | Col-0 | > 40 |  |  |  |  |
|  | *tt4* | > 40 |  |  |  |  |  |  |  |
| DMACA | Col-0 | > 60 |  |  |  |  |  |  |  |
|  | *tt4* | > 40 |  |  |  |  |  |  |  |
| Auramine O | Col-0 | > 60 |  |  |  |  |  |  |  |
|  | *tt4* | > 40 |  |  |  |  |  |  |  |
| CW | Col-0 | > 40 |  |  |  |  |  |  |  |
|  | *tt4* | > 40 |  |  |  |  |  |  |  |

